# Analysis of single nucleotide substitution in CYP21A2 gene using Alphagenome

**DOI:** 10.64898/2026.10.15.759360

**Authors:** Anna Matveeva, Yingtong Xu, Amit V. Pandey

**Author notes:** For correspondence and reprint requests: Prof. Dr. Amit V Pandey, KIKL C837, Pediatric Endocrinology, University Children’s Hospital Bern, Freiburgstrasse 15, 3010 Bern Switzerland. Disclosure Statement: The authors have nothing to disclose.

## Abstract

**Context:** CYP21A2 encodes steroid 21-hydroxylase, which supplies precursors for adrenal glucocorticoid and mineralocorticoid synthesis. Interpretation of its variants is complicated by a homologous pseudogene, complex alleles, heterogeneous functional assays and uncertain noncoding effects. We integrated observed variation with a reference-locus saturation analysis to distinguish transcriptional, splicing and protein-level mechanisms.

**Methods:** The NM_000500.9 MANE transcript was reconstructed from GRCh38 genomic sequence and verified against RefSeq and UniProt P08686. ClinVar, gnomAD and UniProt records were retrieved with provenance, and a cutoff-limited PubMed search was reconciled against Entrez. AlphaGenome Atlas scores were obtained for every possible single nucleotide substitution across the 3,230-bp transcript span. Reported non-SNV alleles were analyzed by direct AlphaGenome inference. Adrenal, testis, ovary, placenta, liver and kidney expression predictions were calculated.

**Results:** The results include 483 ClinVar records, 3,208 gnomAD records and 82 UniProt natural-variants. All 9,690 locus-wide substitutions were scored, comprising 6,018 exonic and 3,672 intronic substitutions. The coding set contained 3,201 missense, 1,098 synonymous, 147 stop-gained, nine start-lost, eight stop-lost and one stop-retained substitution. Direct inference was completed for 413 additional reported alleles and four context controls. Established splice defects, including c.293-13C>G and c.738+75C>T, showed appreciable splicing scores without corresponding large reductions in predicted total RNA. Clinical classification groups differed in AVI score distributions.

**Conclusions:** A sequence-verified CYP21A2 catalogue connects clinical annotations, population observations, splicing predictions and measured enzyme function. Complete single-base coverage of the defined *CYP21A2* transcript was achievable. Regulatory predictions should guide mechanistic experiments and remain separate from enzyme activity, tissue function and clinical classification.

## Introduction

Steroid 21-hydroxylase occupies two central positions in adrenal steroidogenesis. CYP21A2 converts progesterone to 11-deoxycorticosterone and 17-hydroxyprogesterone to 11-deoxycortisol (**Figure 1A**) [1]. Subsequent mitochondrial reactions support mineralocorticoid and glucocorticoid production. The microsomal CYP21A2 enzyme receives reducing equivalents from NADPH through the FAD and FMN cofactors of P450 oxidoreductase (POR) and defects in POR also give CYP21A2 deficiency [2–6]. Its catalytic contribution therefore depends on the amount and integrity of CYP21A2, substrate recognition, heme chemistry, membrane organization and productive electron transfer. Human structural and biochemical work has established a progesterone-bound CYP21A2 structure and connected mutation sites with substrate handling and protein stability [7–9].

**Figure 1:**
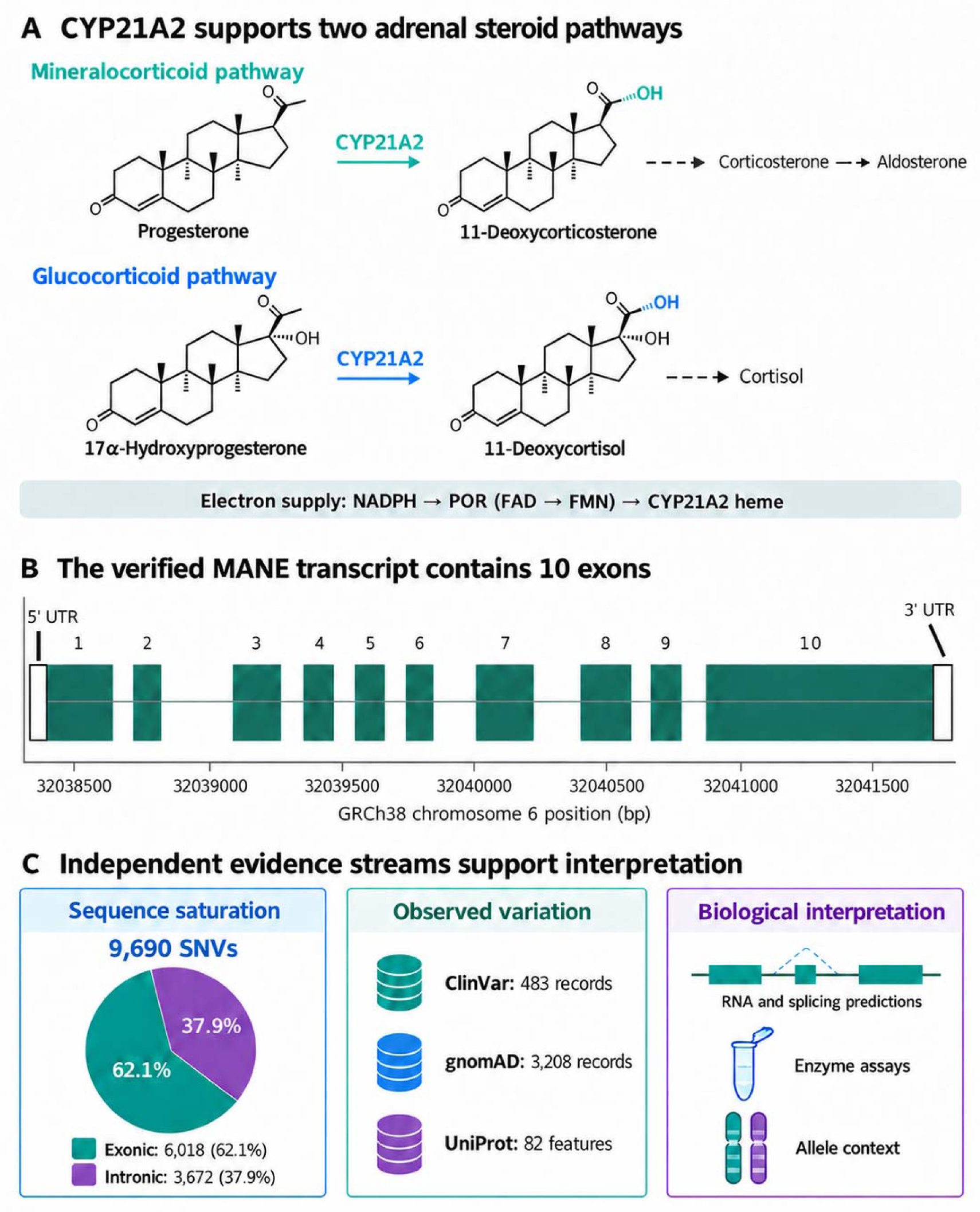
CYP21A2 biology and reference framework. A, The two 21-hydroxylation reactions and downstream steroid products. The electron-transfer annotation depicts NADPH to POR FAD to POR FMN to CYP21A2 heme. B, The ten NM_000500.9 exons on GRCh38; boxes include untranslated sequence and are drawn to genomic scale. C, Evidence streams and saturation denominators. Source counts are not additive because records overlap and differ in type.

Biallelic loss of adequate CYP21A2 function causes 21-hydroxylase-deficient congenital adrenal hyperplasia [10–12]. Reduced cortisol synthesis stimulates ACTH-driven adrenal steroid production, while the extent of mineralocorticoid impairment and diversion toward androgen pathways contributes to the clinical spectrum [13]. Genotype–phenotype associations are informative but incomplete [14, 15]. Large family and patient cohorts demonstrate both recurring severe alleles and variability among milder genotypes [16, 17]. A quantitative assay performed under one expression system cannot be translated directly into an individual patient’s steroid output or prognosis [18].

The genomic setting is unusually difficult. CYP21A2 lies in the RCCX region beside the highly homologous CYP21A1P pseudogene. Gene conversion, deletions, duplications and CYP21A1P/CYP21A2 chimeras can transfer multiple changes together or alter the number of functional copies [19, 20]. Conventional short-read observations may not resolve the gene of origin or allele phase. Long-read studies have demonstrated the utility of locus-specific sequencing for identifying complex alleles and reconciling copy number with sequence changes [21–24]. The distinction between a variant record and a disease-causing haplotype is essential: a truncating change in one copy need not abolish function of an allele carrying another intact CYP21A2 copy [25, 26].

Coding variants have received extensive functional attention, but regulatory and intronic mechanisms remain less evenly characterized [27, 28]. The recurrent intron-2 splice defect was established by early expression studies, while more recent reports describe canonical acceptor changes, cryptic splice-site activation and deep intronic retention [29–32]. Promoter conversion can modify the phenotype associated with an apparently mild coding variant, emphasizing the limits of coding-only interpretation [19, 33, 34]. Historical mutation catalogues provide a valuable foundation, but their totals reflect their date, sources and variant definitions [35].

AlphaGenome predicts multiple molecular readouts from DNA sequence, including gene expression and splicing. Its Atlas provides precomputed scores that permit dense interrogation of a defined locus. The AlphaGenome Variant Impact score, AVI, integrates regulatory features with protein-related and conservation features. These resources make it possible to evaluate all single-base substitutions, while preserving the distinction between a model prediction and an experimentally measured change [36]. The Atlas report was a preprint at the present cutoff. Here we construct a reproducible CYP21A2 catalogue, examine observed and hypothetical variation separately, and use tissue-resolved predictions and published assays to identify mechanistic priorities.

## Methods

### Source data

Supplementary Data 1 is the variant catalogue workbook; Supplementary Data 2 contains the full tabular outputs; Supplementary Data 3 contains the unaggregated prediction scores; and Supplementary Data 4 contains the source records and analysis code in two archive parts. Their contents are described in the Supplementary data file descriptions.

### Reference sequence and scope

The analysis was anchored to GRCh38 chromosome 6, accession NC_000006.12, positions 32,038,415–32,041,644 inclusive, on the positive strand. The MANE Select transcript NM_000500.9 and corresponding NP_000491.4 protein were used for transcript and protein numbering. The genomic interval was retrieved in GenBank format with sufficient local flanking sequence. Concatenation of its ten annotated exons exactly reproduced the 2,006-nt RefSeq transcript. Translation of the 1,488-nt coding sequence, including the termination codon, reproduced the 495-residue UniProt P08686 sequence. Coordinate checks used explicit conversion between one-based inclusive genomic positions and zero-based half-open model intervals. Exon coordinates are provided in Supplementary Data 1, sheet “Reference exons”, and Supplementary Data 2, exon_reference.tsv. The genomic and transcript GenBank records, gene metadata and source protein sequence are retained in Supplementary Data 4.

At every nucleotide in the 3,230-bp MANE span, the reference base was replaced in turn by each of the other three canonical DNA bases. This generated a finite set of 9,690 single nucleotide variants (SNVs). Exonic coverage includes both untranslated regions. Coding consequences were calculated by direct codon substitution and translation, with separate labels for synonymous, missense, stop gain, start loss, stop loss and stop retention. These labels describe sequence consequences; start-loss annotation does not establish whether alternative translation initiation occurs. Intronic positions were classified as canonical splice positions at distance 1–2 from an exon boundary, extended splice region at distance 3–8, or other intronic positions beyond eight bases. These are operational location bins, not pathogenicity thresholds. Every reference base in the saturation set was checked against the retrieved GRCh38 sequence. The complete 9,690-SNV annotation and prediction table is supplied as all_locus_SNV_predictions.tsv in Supplementary Data 2. Its exonic and intronic subsets are accessible in Supplementary Data 1, sheets “Exonic SNVs” and “Intronic SNVs”; field definitions and scope notes are provided in “Dictionary” and “Overview”.

All possible exonic variants was operationalized as all possible single-base substitutions in the selected transcript. Arbitrarily long insertions, deletions, rearrangements and combinations do not form an equivalently bounded set. Reported non-SNV alleles were retained separately, and the analysis did not simulate every possible amino-acid replacement requiring multiple nucleotide changes. Alternative transcripts, extended promoters and structural haplotypes require separate denominators.

For multi-base VCF alleles, compact region summaries use the anchor coordinate. Alleles spanning a boundary can affect more than one region, so the original MANE consequence and complete allele sequence are retained for interpretation. Equivalent indel representations were not collapsed across sources without a validated normalization step.

### Clinical population and protein database retrieval

ClinVar was searched through NCBI Entrez using CYP21A2[gene]. All 483 returned variation identifiers were retrieved as summaries and full VCV XML records. Preserved fields include accession version, clinical classification, review status, evaluation and update dates, conditions, genes, MANE HGVS, GRCh38 allele representation, dbSNP identifiers and PubMed citations. The preliminary Clinical Tables search returned 420 records and was not used as the final completeness denominator. The gene-indexed ClinVar set includes complex and multigene events; it is not a set of 483 independent single-site alleles. The clinical catalogue and intronic subset are provided in Supplementary Data 1, sheets “ClinVar” and “Clinical intronic”, and Supplementary Data 2, clinvar_catalogue.tsv and clinvar_intronic.tsv. The Entrez search, retrieval manifest, summaries and VCV XML records are preserved in Supplementary Data 4.

The gnomAD GraphQL gene endpoint was queried for CYP21A2 on GRCh38 with dataset identifier gnomad_r4. All 3,208 returned records were retained with exome and genome allele counts, allele numbers, allele frequencies, filter labels and source consequence. Missing exome or genome observations were kept missing. Frequencies were not averaged across sequencing datasets, and low-confidence or filtered records were not silently removed. The API gene set spans annotations beyond the selected MANE transcript, so out-of-scope positions were retained and labeled. These data document observed alleles in a population resource; they neither constitute an unselected clinical cohort nor guarantee accurate paralogue assignment at every site. Population observations are provided in Supplementary Data 1, sheet “gnomAD”, and Supplementary Data 2, gnomad_variants.tsv and gnomad_intronic.tsv; the retrieved population payload and filter metadata are retained in Supplementary Data 4.

UniProt P08686 was retrieved as structured JSON. All 82 natural-variant features were extracted with residue coordinates, reference and alternative sequences, descriptions and evidence citations. Protein features were kept as their own evidence unit. Single-base nucleotide routes were linked only when sequence translation produced the same reference and alternative amino acid. Multiple equivalent nucleotide routes were preserved rather than choosing one without source evidence. The original features and nucleotide mappings are provided in Supplementary Data 2, uniprot_variants.tsv and uniprot_variants_mapped.tsv, with the mapped records also presented in Supplementary Data 1, sheet “UniProt”. The retrieved UniProt payload is retained in Supplementary Data 4.

### Literature retrival and functional evidence

The PubMed query was (CYP21A2 OR CYP21B OR “21-hydroxylase”) AND (variant* OR mutation* OR intron* OR splic*), with a publication-date cutoff of 1 October 2026. Complete connector pagination initially returned 1,533 verified articles from a reported 1,539 hits. Entrez reconciliation recovered six book-chapter records. Search results were retained for audit, while selected primary studies were examined for intronic mechanisms, measured activity, structural variation and tissue expression. The retrieval was systematic at the search-inventory level; it was not duplicate independent full-text screening of every record. It therefore does not establish extraction of every published CYP21A2 allele. The reconciled record inventory is provided in Supplementary Data 1, sheet “PubMed inventory”, and Supplementary Data 2, PubMed_inventory.tsv; PubMed_article_inventory.ris contains the 1,532 journal-article records, excluding the six book records. Search reconciliation and curated bibliographic records are retained in Supplementary Data 4.

A 103-row published functional compilation was recovered from the supplementary material of Prado and colleagues, retaining original genomic coordinates, phenotype group, substrate activities, reported standard deviations and cited assay references [37]. All 103 source genomic substitutions matched the current saturation set and encoded their listed protein substitutions. Source-quality issues were retained, including an ambiguous activity cell, a negative background-corrected value and a discrepancy between the stated and tabulated number of progesterone measurements. The compilation was not treated as 103 newly revalidated independent experiments. A second evidence set comprised the approximate Table 1 values for 11 variants in our recent study, with progesterone activity, 17-hydroxyprogesterone activity and protein abundance [38]. The two studies were kept separate; incompatible normalization schemes were not pooled. The two assay datasets are provided separately in Supplementary Data 1, sheets “Assays 103” and “Assays 11”, and Supplementary Data 2, published_103_assay_records.tsv and experimental_function_benchmark.tsv. Their extracted source tables and curation records are preserved in Supplementary Data 4.

**Table 1:** Scope and data coverage. Counts across sources overlap and must not be added to estimate unique observed variants. Electronic scope summaries are provided in Supplementary Data 1, sheets “Overview” and “Main tables”, and Supplementary Data 2, statistics.json and main_tables.json.

| Evidence set | Records or variants | Interpretation |
| --- | --- | --- |
| ClinVar gene-indexed records | 483 | Includes small variants, complex alleles and multigene events |
| ClinVar mapped GRCh38 alleles | 425 | Coordinate-linked; not all lie within the MANE span |
| gnomAD gene-query records | 3,208 | Exome and genome observations retained separately |
| UniProt natural-variant features | 82 | Protein-level evidence, not a genomic denominator |
| PubMed cutoff-limited inventory | 1,538 | Search inventory; selected full-text and evidence extraction |
| All MANE-span SNVs | 9,690 | Complete three-alternative-base substitution set |
| All exonic SNVs | 6,018 | Includes coding region and both UTRs |
| All intronic SNVs | 3,672 | Includes 1,119 observed in ClinVar or gnomAD |
| Reported additional non-SNV alleles | 413 | Direct 16 kb inference; distinct from Atlas scores |
| Published assay records | 103 + 11 | Separate source studies; no pooled activity estimate |

### AlphaGenome Atlas and direct inference

The AlphaGenome Python client version 0.9.0 was used with the user-authorized API account. The service-provided human metadata were archived before scoring. Atlas queries covered the complete MANE genomic span and requested AVI_SCORE, RNA_SEQ, SPLICE_SITE_USAGE, SPLICE_SITES and SPLICE_JUNCTIONS, filtered to CYP21A2 for gene-centric outputs. Six tissue ontology terms were retained: adrenal gland UBERON:0002369, testis UBERON:0000473, ovary UBERON:0000992, placenta UBERON:0001987, liver UBERON:0002107 and kidney UBERON:0002113. Available tracks, assay types, developmental labels, data sources and ontology identifiers were preserved. The retrieved Atlas gene identifier was ENSG00000231852.9. MANE-based variant annotation and the model’s gene masks are distinct annotation layers. The 48 scorer–track metadata records are provided in Supplementary Data 1, sheet “Track metadata”, and Supplementary Data 2, track_metadata.tsv. Service metadata objects are retained in Supplementary Data 4.

Atlas supplied 15 tissue tracks for each of RNA, splice-site usage and splice-junction scoring, plus two tissue-independent splice-site classes and one AVI score. Queries were checkpointed in bounded genomic chunks. All 9,690 expected SNVs were returned and reference-validated. The AVI model features and feature attributions were additionally retrieved for every SNV, including the AlphaMissense feature. AVI raw values, server-calibrated values and molecular scores were stored separately. AVI is a ranking score; neither its raw value nor the returned calibrated quantile is a patient-level probability of disease. Feature attribution describes the composite model’s calculation and does not independently validate a biological mechanism. All SNV-level model features and attributions are supplied in Supplementary Data 2 as AVI_SCORE_MODEL_FEATURES.tsv and AVI_SCORE_FEATURE_IMPORTANCE.tsv. Full AVI outputs are available in Supplementary Data 3 as raw_AVI_SCORE.tsv.gz; the cached Atlas and feature-query objects are retained in Supplementary Data 4.

The Atlas API rejected non-SNV queries with an explicit unsupported-allele message. A direct 1,048,576-bp inference request exceeded the account’s per-minute prediction-token quota. Consequently, 413 distinct reported non-SNV alleles within the MANE span were analyzed with direct AlphaGenome inference in 16,384-bp variant-centered windows. Four established splice-control SNVs were run through the same protocol to document context sensitivity. All 417 direct requests were completed after retrying transient quota failures. This shorter-context analysis is clearly labeled throughout the files and is not presented as interchangeable with the Atlas protocol. Direct inference supplied molecular scores but no AVI score; missing AVI values were retained as unavailable. The 417 direct-inference summaries and four paired context controls are provided in Supplementary Data 1, sheets “Live 16kb” and “Context controls”, and Supplementary Data 2, live_16kb_predictions.tsv and context_sensitivity_controls.tsv. Completion status is recorded in live_scoring_status.tsv. The full direct outputs are supplied in Supplementary Data 3 as raw_live_16kb.tsv.gz, with individual cached results and the input allele list in Supplementary Data 4.

### Aggregation descriptive comparisons and validation

For a given tissue, the signed RNA score with the largest absolute value across available tracks was retained as a compact summary. For nonnegative splice-site usage and splice-junction scores, the maximum across relevant tracks was used; junction summaries also maximized across returned junctions. Complete per-track and per-junction matrices remain available. These maxima summarize a model response, not mean tissue expression or a biological replicate estimate. The score summary can be sensitive to the number and type of available tracks, particularly for placenta and other tissues lacking a GTEx counterpart. Unaggregated Atlas outputs are provided in Supplementary Data 3 as raw_RNA_SEQ.tsv.gz, raw_SPLICE_SITE_USAGE.tsv.gz, raw_SPLICE_SITES.tsv.gz and raw_SPLICE_JUNCTIONS.tsv.gz. These retain the track-specific and, where applicable, junction-specific information underlying the compact tissue summaries.

Clinical classifications were grouped as pathogenic or likely pathogenic, uncertain significance, benign or likely benign, conflicting, or other. Score distributions were compared descriptively among mapped SNVs. No clinical decision threshold, sensitivity, specificity or calibrated pathogenicity model was derived. Shared ascertainment, related alleles, evidence reuse and possible training overlap prevent interpretation as independent clinical validation. Tissue correlations were calculated using Spearman rank correlation across the complete saturation set and are descriptive properties of the predictions. No inferential P values were assigned to correlated substitutions. Known intronic variants were distinguished from hypothetical variants by exact matches to retrieved ClinVar or gnomAD alleles, supplemented by curated literature examples.

Quality controls included expected-versus-observed SNV counts, uniqueness of chromosome-position-reference-alternate keys, genomic reference checks, transcript reconstruction, protein sequence identity, source record reconciliation and separation of complex alleles. Summary counts and selected descriptive statistics are provided in Supplementary Data 2, statistics.json. Supplementary Data 4 includes the analysis scripts, dependency list, README and SHA256SUMS.txt file; the README documents the scope of the archived workflow and the distinction between cached outputs and future API reruns.

## Results

### Reference validation and source coverage

The verified transcript comprised 2,006 exonic nucleotides and 1,224 intronic nucleotides across ten exons and nine introns (**Figure 1B**). The coding region contained 1,485 translated nucleotides plus a three-base termination codon; the selected transcript additionally contained an eight-base 5′ untranslated region and a 510-base 3′ untranslated region. This transcript-specific boundary definition explains why some database variants labeled untranslated fall outside the present MANE span. All reconstructed exon sequences and the translated protein matched the respective reference records exactly.

ClinVar contributed 483 gene-indexed records, of which 425 had explicit GRCh38 VCF alleles and 420 had NM_000500.9 HGVS annotations (**Figure 2A and B**). The complete record set included 164 pathogenic or likely pathogenic, 160 uncertain, 116 benign or likely benign, 31 conflicting and 12 other classifications (**Figure 2C**). MANE-based coordinate annotation assigned 301 records to coding sequence, 83 to introns, 23 to the 3′ untranslated region and two to the 5′ untranslated region; 16 lay outside the selected span and 58 were complex or not represented as one mapped allele (**Figure 2D**). These categories describe record assignment, not unique patients or independent disease-causing haplotypes

**Figure 2:**
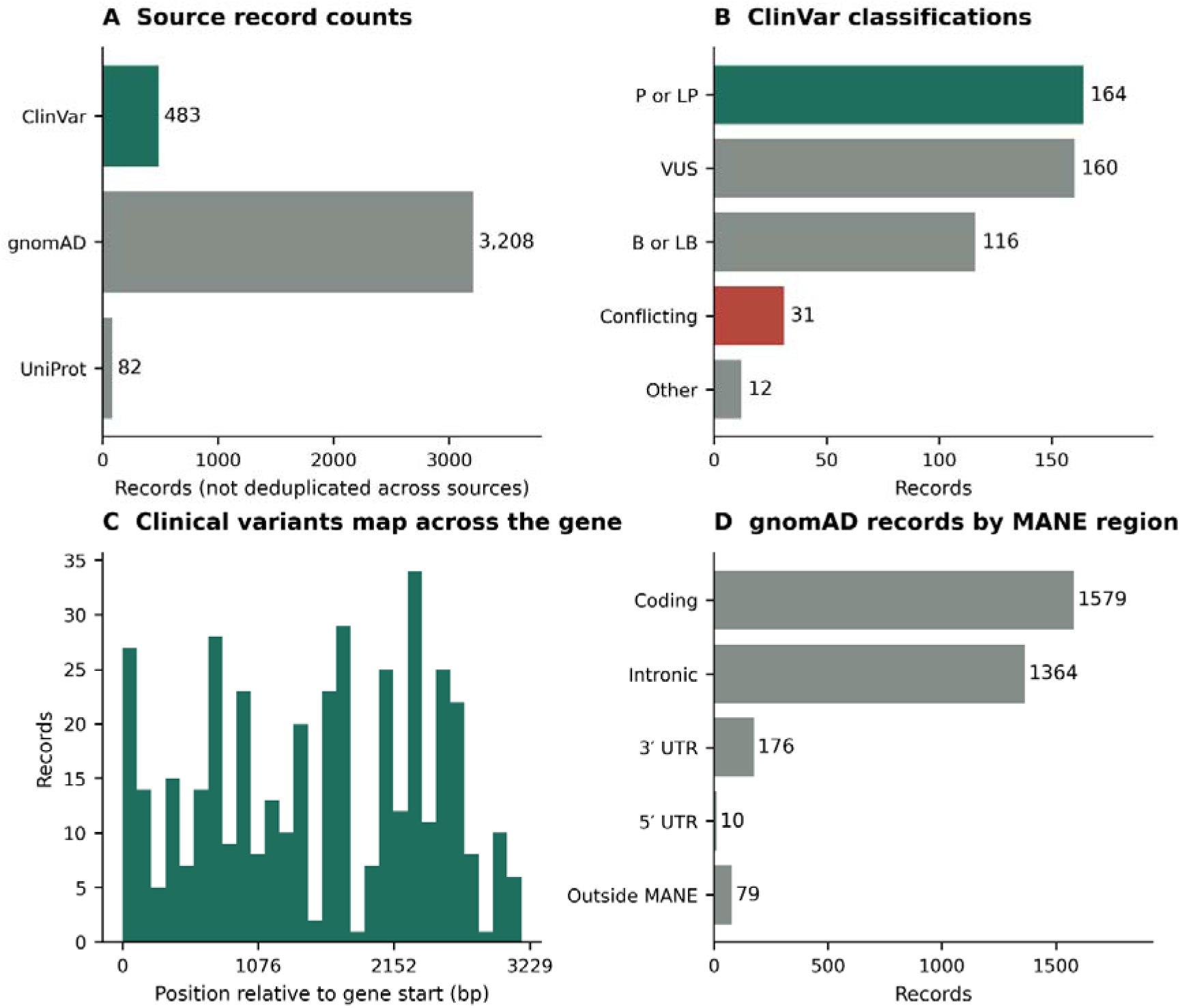
Source coverage and clinical annotations. A, Retrieved ClinVar records, gnomAD records and UniProt natural-variant features. B, Clinical classification groups among all 483 ClinVar records. P or LP, pathogenic or likely pathogenic; B or LB, benign or likely benign; VUS, uncertain significance. C, Genomic distribution of coordinate-mapped clinical records within the MANE span. D, gnomAD records assigned to MANE coordinate regions. These are record counts, not patient counts or estimates of disease prevalence.

The gnomAD query returned 3,208 records (**Figure 2A**), including 3,129 within the MANE span and 79 outside it. Within the span, 1,579 records were assigned to coding sequence, 1,364 to introns, 176 to the 3′ untranslated region and ten to the 5′ untranslated region. Exactly 2,749 gnomAD records matched saturation SNVs. The union of database-observed intronic SNVs contained 1,119 variants, including 74 observed in ClinVar and 1,102 in gnomAD; 57 were shared. The remaining intronic saturation variants are hypothetical with respect to these two retrieved sources. UniProt contributed 82 protein-level features, which were retained independently to avoid mixing protein features and genomic allele counts. The 1,119 database-observed intronic SNVs are enumerated in Supplementary Data 1, sheet “Observed intronic SNVs”, and Supplementary Data 2, known_intronic_SNV_predictions.tsv, with separate ClinVar and gnomAD observation flags.

### The complete exonic substitution landscape

All 6,018 exonic substitutions received Atlas scores. Of these, 4,464 affected the coding sequence, including the termination codon; 1,530 affected the 3′ untranslated region and 24 affected the 5′ untranslated region. Codon translation yielded 3,201 missense, 1,098 synonymous, 147 stop-gained, nine start-lost, eight stop-lost and one stop-retained consequence. The full nucleotide-level catalogue provides reference and alternative codons, amino-acid changes, genomic identifiers and tissue scores. For the protein-level projection, blank amino-acid cells represent substitutions inaccessible by one nucleotide change, rather than missing predictions (**Figure 3A**). All 6,018 exonic substitutions are listed in Supplementary Data 1, sheet “Exonic SNVs”, and Supplementary Data 2, all_exonic_SNV_predictions.tsv. The TSV includes the fuller sequence annotation, including codon fields, whereas the workbook presents selected annotation and score columns.

**Figure 3:**
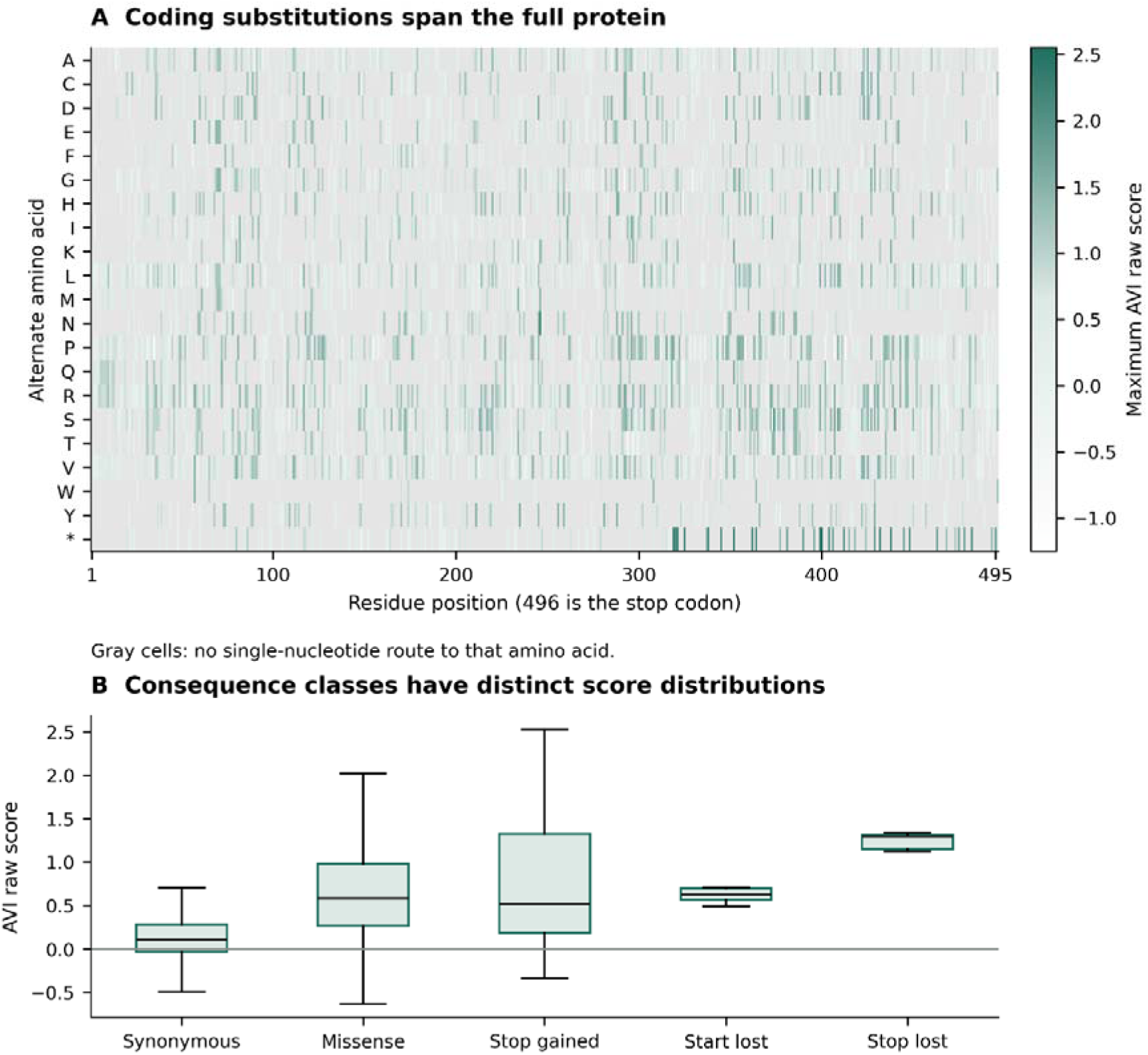
Exonic saturation landscape. A, Maximum AVI raw score for each amino-acid substitution reachable through a single nucleotide change. The horizontal axis uses the 495-residue protein plus termination codon at position 496; gray cells lack a single-base route. When more than one SNV encodes a substitution, the maximum is shown only for visualization and the individual scores remain in the data. B, AVI distributions across sequence-consequence classes. Boxes show quartiles and median; whiskers extend to 1.5 interquartile ranges and outliers are omitted from display. Scores are deterministic model outputs and are not enzyme activities or probabilities of pathogenicity.

The model’s response was heterogeneous within consequence classes (**Figure 3B**). Protein-disrupting categories did not imply large predicted splicing or RNA changes. For example, c.518T>A (p.Ile173Asn) had an AVI raw score of 1.225 with an adrenal splice-usage score of 0.0078, whereas c.955C>T (p.Gln319Ter) had an AVI raw score of 2.247 and an adrenal splice-usage score of 0.0156. Such differences are consistent with a composite impact score incorporating protein-level features. They do not show that either allele preserves enzyme activity. Representative clinically important coding substitutions are listed in Table 2.

**Table 2:** Representative coding substitutions and adrenal scores. Protein labels use one-letter amino-acid notation. RNA is the selected signed gene-expression log-fold-change score; usage is a nonnegative splicing effect score. These are predictions, not residual enzyme activities. c.844G>C also encodes Val282Leu, and c.1179C>G also encodes His393Gln; each nucleotide allele has its own score. Individual nucleotide-allele scores are available in Supplementary Data 1, sheets “Exonic SNVs” and “ClinVar”, and Supplementary Data 2, all_exonic_SNV_predictions.tsv; the original table block is retained in “Main tables” and main_tables.json.

|  | Protein | AVI raw | RNA raw | Usage raw |
| --- | --- | --- | --- | --- |
| **MANE cDNA** |  |  |  |  |
| c.92C>T | p.P31L | 0.857 | 0.0248 | 0.0117 |
| c.518T>A | p.I173N | 1.225 | 0.0067 | 0.0078 |
| c.844G>T | p.V282L | 1.019 | -0.0142 | 0.0137 |
| c.955C>T | p.Q319* | 2.247 | 0.0118 | 0.0156 |
| c.1069C>T | p.R357W | 0.531 | -0.0030 | 0.0137 |
| c.1179C>A | p.H393Q | 0.930 | 0.0100 | 0.0156 |
| c.1306C>T | p.R436C | 0.605 | -0.0113 | 0.0078 |
| c.1451G>T | p.R484L | 1.206 | -0.0139 | 0.0078 |

### Intronic variants reveal splicing-specific mechanisms

The median adrenal splice-usage score was 0.4609 for the 108 possible substitutions at canonical splice positions, compared with 0.0508 for 324 extended splice-region substitutions and 0.0117 for 3,240 other intronic substitutions (**Figure 4A**). These distributions support a location-dependent enrichment of model-predicted splice disruption, while leaving clear exceptions away from exon boundaries. The highest adrenal splice-usage scores included substitutions at the intron-5 donor, such as c.651+2T>G. The complete 3,672-SNV intronic prediction set is provided in Supplementary Data 1, sheet “Intronic SNVs”, and as the intronic subset of all_locus_SNV_predictions.tsv in Supplementary Data 2.

**Figure 4:**
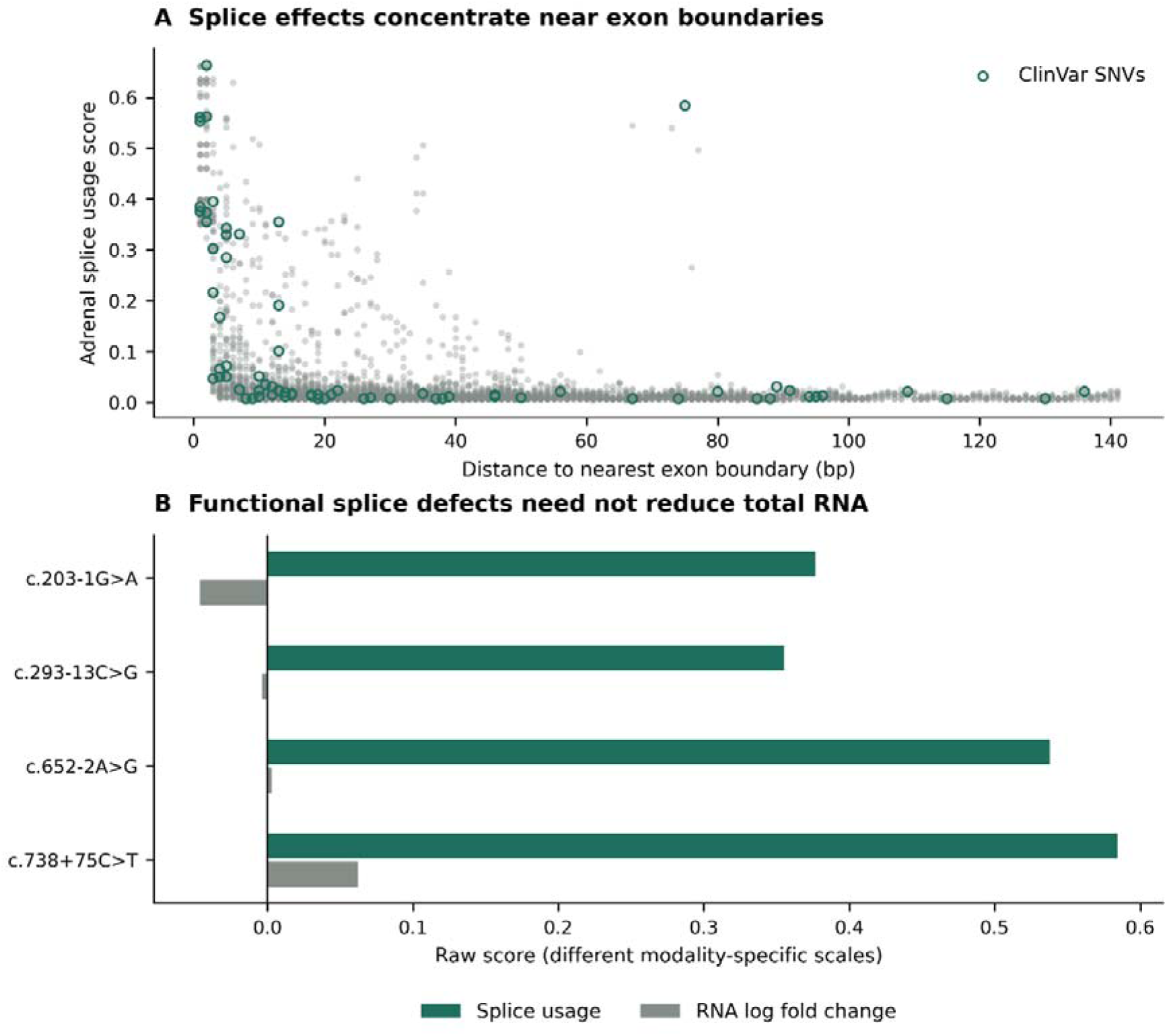
Intronic location and mechanistic controls. A, Adrenal splice-site-usage score versus distance to the nearest exon boundary for all 3,672 intronic SNVs; outlined points are ClinVar-observed SNVs. B, Four splice-control alleles with published functional evidence. The bars display splice-usage and signed RNA scores on their own modality-specific raw scales. Comparing their absolute magnitudes across modalities is not a calibrated effect-size comparison. The panel illustrates that a strong splice response can coexist with a small RNA abundance score.

The recurrent c.293-13C>G allele had an adrenal usage score of 0.3553 and junction score of 6.207, while its predicted RNA score was −0.0038. The deep intronic c.738+75C>T allele had corresponding values of 0.5843, 6.133 and +0.0621 (**Figure 4B and Table 3**)The latter variant has now been associated with 73-base intron-6 retention in a minigene study, whereas an earlier report lacked functional validation [32, 39]. The predicted splice response therefore aligns with the reported mechanism, but the RNA score does not reproduce a simple transcript-loss narrative. The experiments predict a frameshift and likely nonsense-mediated decay; our model results do not establish the magnitude of decay in native adrenal cells.

**Table 3:** Selected intronic variants and evidence. Adrenal raw scores are shown. Legacy IVS2+1 and IVS9-9 labels were mapped by intron position and checked for the stated reference base. The latter is c.1223-9C>A on the selected transcript. Clinical reports and functional experiments are not equivalent evidence classes. The evidence records are supplied in Supplementary Data 1, sheets “Splice controls” and “Clinical intronic”, and Supplementary Data 2, intronic_benchmarks.tsv and clinvar_intronic.tsv; the original table block is retained in “Main tables” and main_tables.json.

| Variant | Usage | Junction | Published evidence | Ref |
| --- | --- | --- | --- | --- |
| c.203-1G>A | 0.377 | 4.250 | Exon 2 skipping | [40] |
| c.292+1G>A | 0.385 | 3.723 | Exon 2 skipping;<br>additional intron<br>retention in a subset | [54] |
| c.292+5G>A | 0.343 | 3.248 | Severe cis allele<br>reported with<br>Val282Leu | [55] |
| c.293-13C>G | 0.355 | 6.207 | Aberrant intron 2<br>splicing | [29] |
| c.652-2A>G | 0.538 | 5.613 | Cryptic acceptor; 16<br>nt exon deletion | [31] |
| c.738+75C>T | 0.584 | 6.133 | 73 bp intron 6<br>retention | [32] |
| c.940-1G>C | 0.461 | 5.941 | Splice acceptor<br>variant reported by<br>long-read genotyping | [21] |
| c.1223-9C>A | 0.351 | 6.250 | Cryptic acceptor at -<br>7; loss of protein and<br>activity | [30] |

**Table 4:**
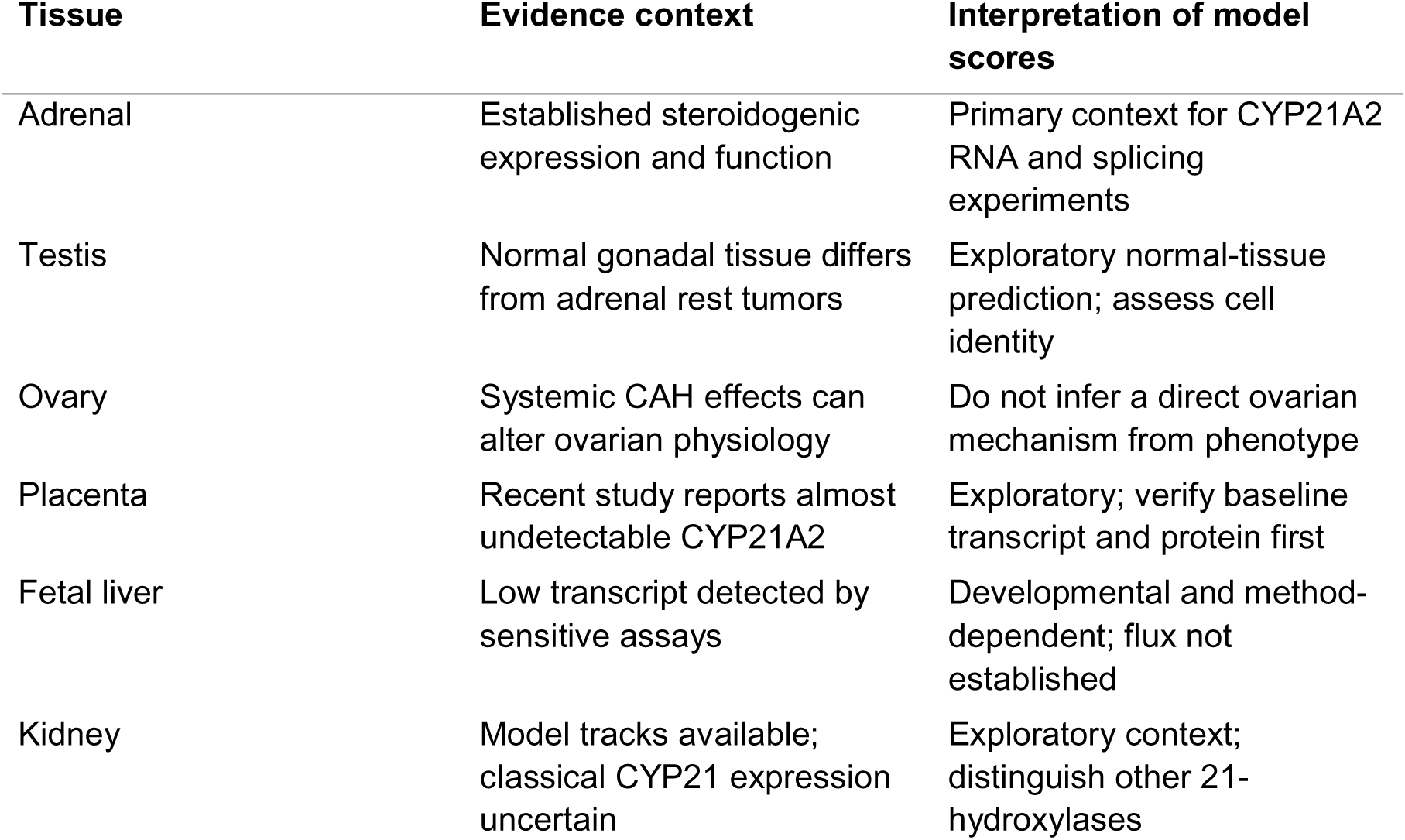
Tissue contexts and permissible interpretation. Testicular adrenal rest tumors express an adrenal-like program and should not be equated with ordinary testis [56]. Ovarian abnormalities in a humanized CYP21A2 mouse model do not by themselves demonstrate a cell-autonomous ovarian enzyme role [57]. The original tissue-interpretation table is retained in Supplementary Data 1, sheet “Main tables”, and Supplementary Data 2, main_tables.json; model-track annotations are supplied separately in “Track metadata” and track_metadata.tsv.

Other mechanism controls included c.203-1G>A, experimentally associated with exon-2 skipping, and c.652-2A>G, which activates a cryptic acceptor and removes 16 nucleotides from exon 6 [31, 40]. Their adrenal usage scores were 0.3768 and 0.5378, respectively (**Table 3**). Reported intronic changes without RNA validation were retained at a lower mechanistic evidence level. In particular, disease association and in silico splice-site prediction were not relabeled as demonstrated exon skipping [41]. Table 3 presents selected evidence controls, while the full clinical intronic table retains every retrieved record and its classification. The eight curated intronic evidence examples are provided in Supplementary Data 1, sheet “Splice controls”, and Supplementary Data 2, intronic_benchmarks.tsv. The 83-record clinical intronic subset is available separately in “Clinical intronic” and clinvar_intronic.tsv; these evidence examples should not be conflated with the four direct-inference context controls.

### Tissue-resolved predictions and biological relevance

All six requested tissue contexts had relevant model tracks, but track presence does not establish appreciable endogenous CYP21A2 expression. The adrenal RNA summary ranged from −0.6966 to +0.1599 across the saturation set. The largest negative RNA predictions across tissues clustered in the distal 3′ untranslated region around c.*487–c.*491 (**Figure 5A**). This is a model-derived candidate regulatory interval; its effect on polyadenylation, transcript stability or overlapping-gene regulation requires direct testing. It should not be assigned a specific mechanism from the RNA score alone.

**Figure 5:**
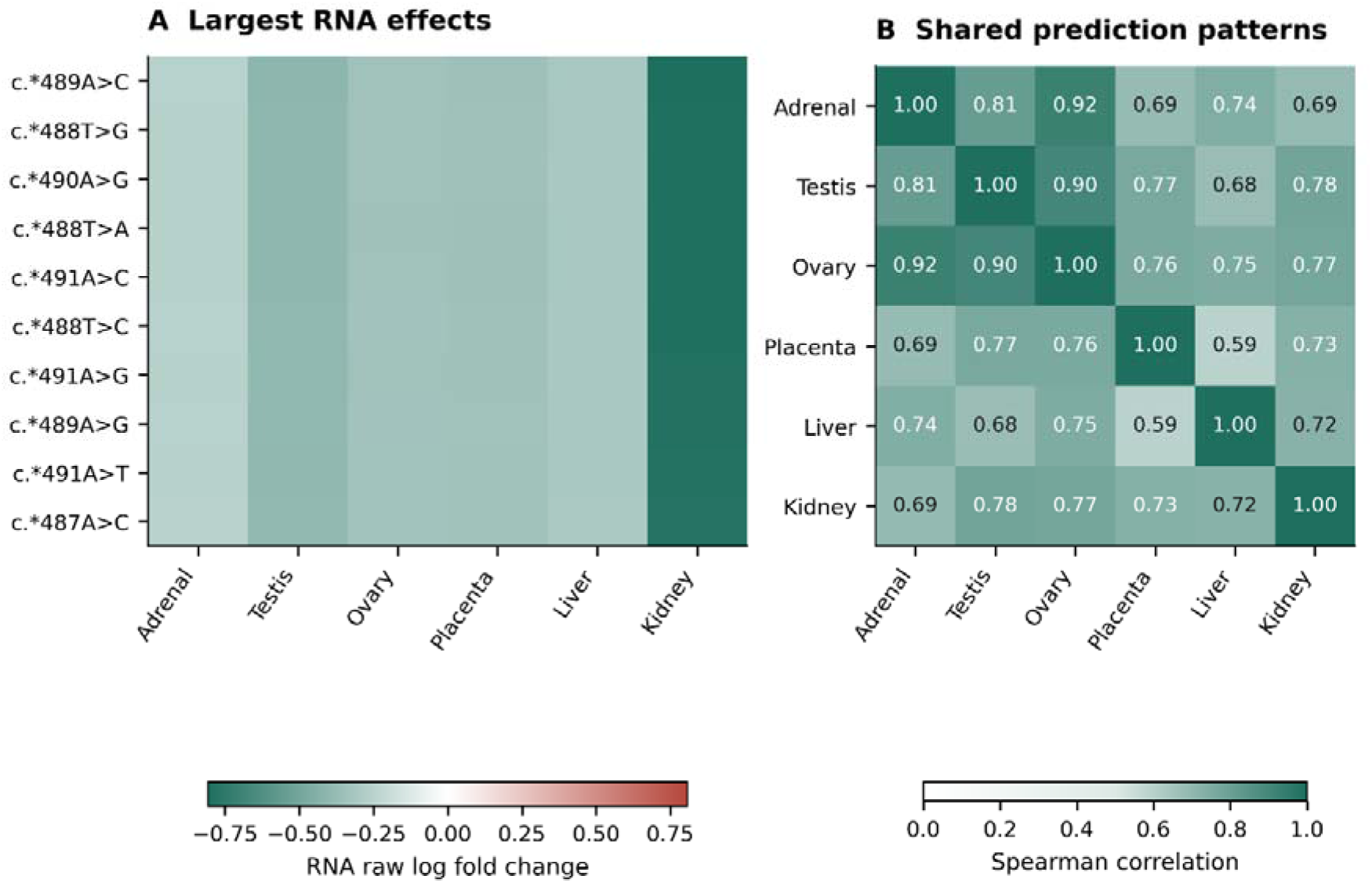
Tissue-resolved RNA predictions. A, Ten substitutions with the largest absolute RNA score across the six tissue summaries, concentrated in the distal 3′ untranslated region. A signed maximum-absolute track score is retained per tissue. B, Spearman correlation across all 9,690 SNVs. These correlations describe model outputs; correlated substitutions are not biological replicates. Track coverage differs by tissue, and a predicted response does not establish endogenous CYP21A2 expression or steroidogenic flux. The underlying six-tissue summaries, full RNA outputs and track annotations are supplied in Supplementary Data 2, all_locus_SNV_predictions.tsv and track_metadata.tsv, and Supplementary Data 3, raw_RNA_SEQ.tsv.gz.

RNA score patterns were correlated across tissues. Adrenal correlations were approximately 0.81 with testis, 0.92 with ovary, 0.69 with placenta, 0.74 with liver and 0.69 with kidney (**Figure 5B**). These correlations describe related model responses to the same substitutions and are not evidence that CYP21A2 contributes equally to steroid flux in those tissues. The placenta had different assay coverage and developmental labels from adult GTEx tissues. Comparisons of maxima across unequal track collections must therefore remain descriptive. The six-tissue summaries underlying Figure 5 are retained in all_locus_SNV_predictions.tsv (Supplementary Data 2), with the corresponding RNA tracks in raw_RNA_SEQ.tsv.gz (Supplementary Data 3) and their assay and tissue annotations in “Track metadata” (Supplementary Data 1).

Experimental tissue evidence identifies the adrenal as the principal physiological context. Fetal adrenal studies support substantial CYP21A2 expression and steroidogenic function [42]. A recent placental study found CYP21A2 almost undetectable, whereas fetal liver work detected low transcript levels using more sensitive methods [43, 44]. Earlier work failed to detect classical CYP21 expression in several extra-adrenal tissues under its assay conditions [45]. These observations are not necessarily contradictory: tissue stage, detection sensitivity and cellular composition differ. Extra-adrenal 21-hydroxylation cannot automatically be attributed to CYP21A2.

### Clinical concordance without diagnostic overclaiming

Within the 361 ClinVar records matched to the Atlas SNV set, median AVI raw scores were 1.134 for pathogenic or likely pathogenic records, 0.575 for uncertain records, 0.451 for conflicting records and 0.017 for benign or likely benign records. The mapped groups comprised 96, 131, 30 and 98 records, respectively, with six additional records in the other category (**Figure 6**). It does not justify using an AVI threshold to reclassify uncertain alleles or to infer clinical sensitivity and specificity. Record-level classifications, prediction status and scores used for these comparisons are available in Supplementary Data 1, sheet “ClinVar”, and Supplementary Data 2, clinvar_catalogue.tsv; the AlphaMissense feature is retained in AVI_SCORE_MODEL_FEATURES.tsv.

**Figure 6:**
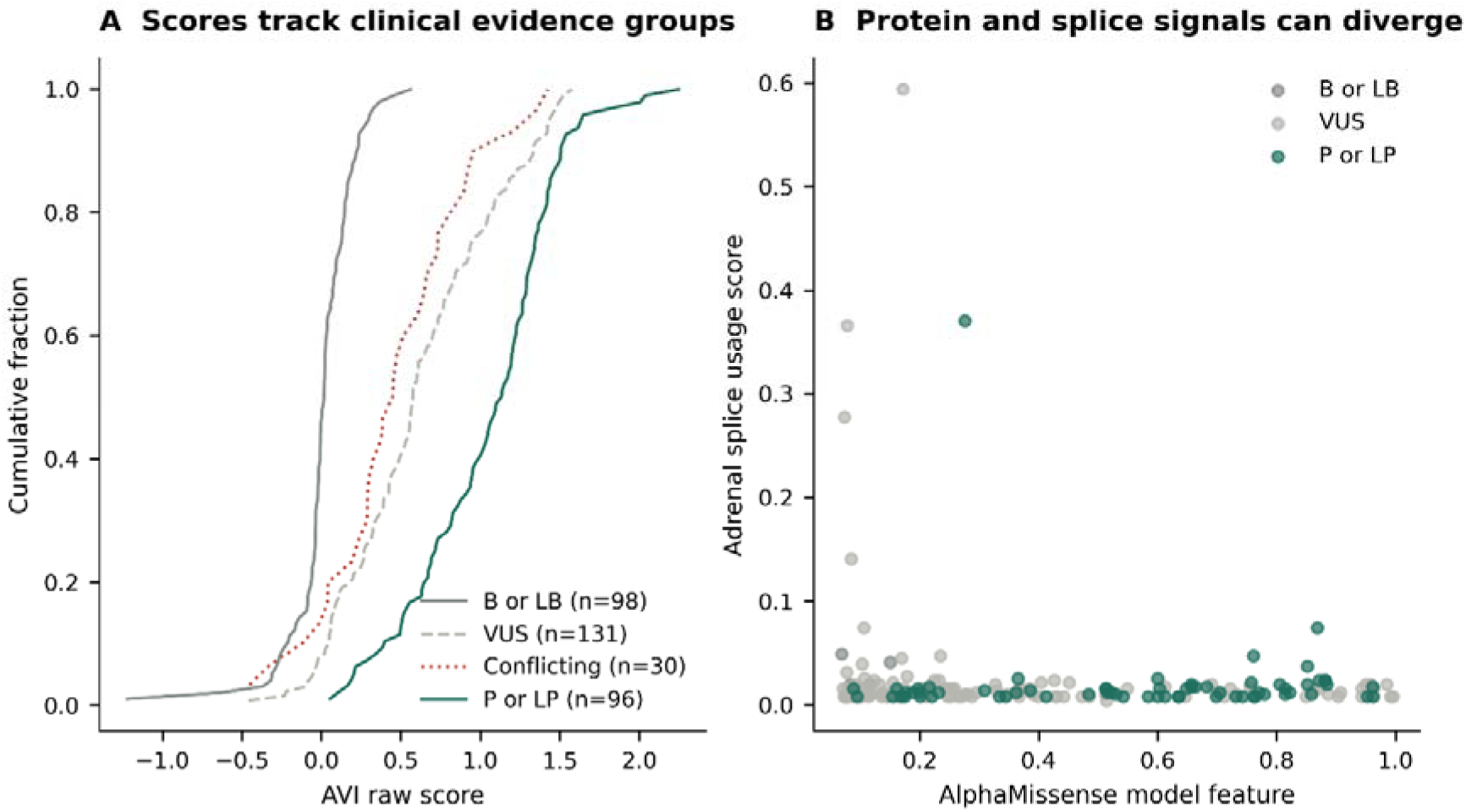
Clinical evidence groups and molecular scores. A, Empirical cumulative AVI distributions for Atlas-matched ClinVar records in four principal classification groups. Six records in other categories are not plotted. B, AlphaMissense feature versus adrenal splice-usage score for matched missense SNVs in the three displayed groups. Clinical records and model features are not independent ground truth; this is a descriptive comparison rather than an accuracy benchmark or basis for reclassification. The underlying clinical records and model features are supplied in Supplementary Data 1, sheet “ClinVar”, and Supplementary Data 2, clinvar_catalogue.tsv and AVI_SCORE_MODEL_FEATURES.tsv.

The additional direct-inference analysis completed all 413 selected reported non-SNV alleles. These include indels and other multi-base allele representations; they are not all independent biological events, because equivalent representations can occur in repetitive sequence. Their scores were recorded in separate live16 columns, with full per-track outputs and four context-control comparisons. The clinical catalogue retains structural or unresolved records even when no single-allele prediction is available. A blank model cell for a complex allele is a scope limitation, not evidence against pathogenicity. Individual non-SNV outputs and paired control comparisons are available in Supplementary Data 1, sheets “Live 16kb” and “Context controls”, with the complete direct-inference matrices in Supplementary Data 3.

### Measured protein function and model interpretation

The 103-row published assay compilation adds an experimental dimension to the coding catalogue [37]. All source genomic SNVs were matched to the current reference. However, assay designs, normalization, substrates and phenotype labels differ across the cited studies [46]. The compilation lists Ile172Asn and Ile173Asn as separate rows; both genomic substitutions translate as labeled on the selected sequence, so adjacency alone does not establish a duplicate. Source records are retained without collapsing them. One ambiguous numerical cell remains missing, and a reported negative activity estimate is preserved rather than forced to zero. These issues preclude treating the compilation as a uniform quantitative training set without further assay-level curation. The source values, missing entries and curation flags remain available in Supplementary Data 1, sheet “Assays 103”, and Supplementary Data 2, published_103_assay_records.tsv.

Our recent 11-variant study illustrates substrate-dependent and abundance-dependent effects (**Figure 7**). Its summary table gives approximate progesterone/17-hydroxyprogesterone activities of 111/99% for Leu10del, 72/77% for Arg76Lys, 58/53% for Glu162Gly, 76/46% for Ser274Tyr, 30/18% for Leu308Val, 92/93% for Ser373Asn, 53/19% for Pro387Leu, 71/44% for His393Gln, 40/41% for Arg401Gly, 26/18% for Arg436Cys and 100/67% for Ser494Asn [38]. These are study-specific protein-normalized activity estimates, not patient-level enzyme function. Protein abundance is a separate measurement. Where two SNVs encode the same amino-acid substitution, a protein label alone does not identify the experimentally tested nucleotide allele. The separate activity, protein-abundance and nucleotide-mapping fields are provided in Supplementary Data 1, sheet “Assays 11”, and Supplementary Data 2, experimental_function_benchmark.tsv.

**Figure 7:**
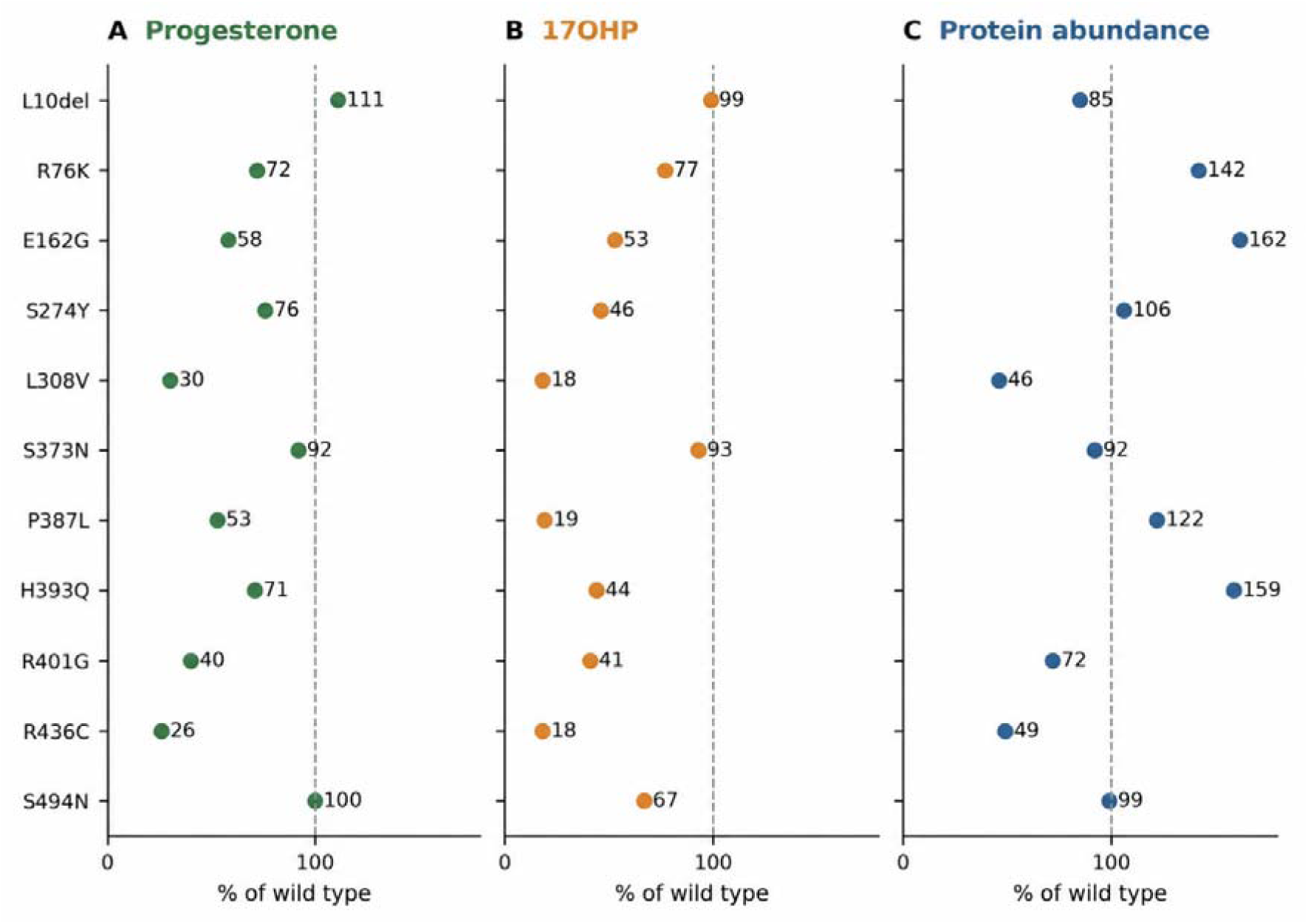
Measured activity and abundance in the recent 11-variant study. A, Progesterone 21-hydroxylation. B, 17-hydroxyprogesterone (17OHP) 21-hydroxylation. C, CYP21A2 protein abundance. Values reproduce the approximate rounded means in Xu et al. Table 1, with wild type set to 100%. Activities are reported as CYP21A2-protein-normalized in that source. Table-specific standard deviations were unavailable and are not invented. The dashed line marks wild type. These study results are not pooled with earlier assays; His393Gln has a substantial cross-study discrepancy. The source values and mapping notes are supplied in Supplementary Data 1, sheet “Assays 11”, and Supplementary Data 2, experimental_function_benchmark.tsv.

His393Gln highlights a material evidence conflict. An earlier primary study reported approximately 2.2% progesterone and 2.5% 17-hydroxyprogesterone activity in a COS-7 system, whereas the recent HEK293(T) study reported approximately 71% and 44% [38, 47]. The earlier assay corrected for total protein, while the later study normalized activity to CYP21A2 abundance. Differences in cellular expression and denominator therefore matter, although they do not fully resolve the discrepancy without reanalysis of the experiments. We retain both study results and do not average them. The Leu10del protein label and its reported cDNA alias also require independent reconciliation before a DNA-specific prediction can be attached.

AVI feature attributions distinguished regulatory and protein-related contributions among representative alleles (**Figure 8A**). Strong splicing-feature contributions were apparent for established intronic splice defects, whereas AlphaMissense and termination features contributed to selected coding variants. Conservation also contributed across classes. These outputs support mechanism-specific follow-up, but a model explanation is not evidence that an experimentally proposed structural pathway is correct. Complete feature values and attributions for all 9,690 SNVs are supplied in Supplementary Data 2, AVI_SCORE_MODEL_FEATURES.tsv and AVI_SCORE_FEATURE_IMPORTANCE.tsv, including the values underlying Figure 8.

**Figure 8:**
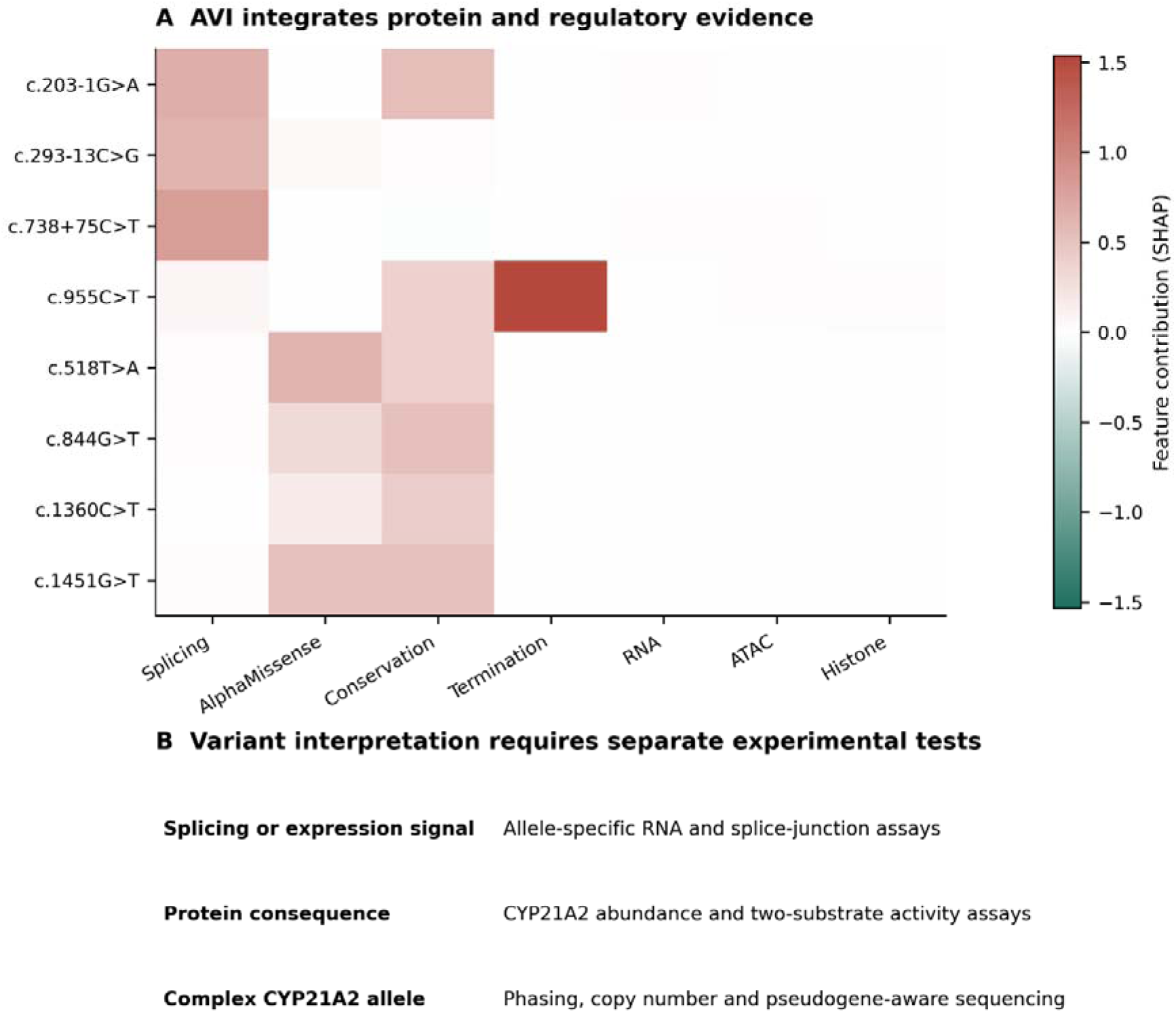
Model components and experimental priorities. A, Selected AVI feature attributions for eight representative nucleotide substitutions. Positive and negative contributions refer to the composite model output. Columns are selected mechanistic features rather than all 18 model features; full values are supplied. The attribution is an explanation of the model computation, not experimental proof of mechanism. B, Follow-up assay classes corresponding to RNA, protein and complex-allele hypotheses. All model feature values and attributions are supplied in Supplementary Data 2, AVI_SCORE_MODEL_FEATURES.tsv and AVI_SCORE_FEATURE_IMPORTANCE.tsv.

## Discussion

This study provides complete single-base coverage of a sequence-verified CYP21A2 transcript span and integrates it with current clinical, population and protein evidence. The resulting resource separates three questions that are often conflated: whether an allele has been observed, whether a model predicts a molecular effect, and whether an experiment or clinical assessment demonstrates disease relevance. The exact substitution denominator makes completeness auditable. By contrast, a statement that every known CYP21A2 mutation has been captured would require broader resources, full-text variant extraction, structural-allele resolution and ongoing updates. The present source-specific counts and unresolved records make that distinction explicit.

The difference between the preliminary 420-record Clinical Tables result and the 483-record primary ClinVar retrieval illustrates why database access routes matter. Likewise, a gene-query population set contains variants beyond the selected transcript, and protein features are not directly interchangeable with nucleotide alleles. Reconciliation should preserve accession versions, genomic assembly and original nomenclature. A simple list of protein names cannot support a reproducible regulatory prediction because distinct nucleotide substitutions can produce the same amino-acid change.

The most direct biological contribution of AlphaGenome in this setting is the opportunity to identify candidate RNA mechanisms for intronic, synonymous and some missense alleles. Strong signals at canonical splice positions are expected, but the deep intronic c.738+75C>T example demonstrates the importance of evaluating beyond a narrow splice-boundary window. The concordance with reported intron retention is useful for prioritization. It does not prove the same transcript abundance, isoform ratio or decay pathway in adrenal tissue. Minigene assays themselves remove parts of the native genomic context and may not capture developmental or cell-state regulation.

Several established splice alleles produced small total-RNA predictions despite appreciable usage and junction scores. This is not a reason to discard the splicing signal. Total abundance and isoform composition are different quantities, and disruption of productive splicing may be buffered, incompletely represented in a model or dependent on RNA surveillance. Conversely, a measurable RNA score without a demonstrated transcript change does not establish reduced enzyme function. Follow-up should therefore quantify both correctly spliced RNA and aberrant products, ideally with allele-specific assays and measurement of endogenous protein.

CYP21A2 deficiency can arise from altered folding, heme incorporation, substrate positioning, membrane association or electron transfer without a large change in transcription. This explains why a low regulatory score cannot rescue a clinically established coding allele. The addition of an AlphaMissense feature to AVI broadens the composite signal, but it does not replace substrate-specific kinetics or cellular activity assays. Earlier human structural and functional studies show that the same global protein can be perturbed through multiple mechanisms [8, 48, 49].

The activity evidence also argues against a universal conversion from percent-of-wild-type activity to phenotype. Protein-normalized catalytic activity and total cellular steroid production answer different questions. A variant can be catalytically competent per molecule yet poorly expressed, or abundant but inefficient. The H393Q discrepancy demonstrates why source assay, normalization and uncertainty must accompany every numerical activity value. Reanalysis should use raw conversion data, expression measurements, substrate concentration and incubation conditions before comparing studies quantitatively. Discordant results are an experimental priority rather than a rationale to select the preferred number.

The reference-locus SNV model cannot reproduce every RCCX haplotype. Gene conversions often transfer several changes in cis; the effects of such combinations need not equal the sum of individually scored substitutions. Recent functional work shows greater impairment for some cis combinations than for their component variants [32]. Copy number can also change interpretation, as illustrated by Gln319Ter on a duplicated allele [25]. The pathogenicity of a sequence change, the functionality of the whole chromosome and the patient’s biallelic genotype are therefore distinct levels of analysis.

Promoter conversion deserves explicit consideration even though exhaustive promoter saturation was outside the present scope. A recent diagnostic series identified promoter variants in 89 of 1,279 tested individuals and in 12.1% of 207 affected patients, reinforcing the value of promoter analysis when a coding-only result does not explain severity [19]. Long-read population screening also found substantial duplication and rare-allele diversity, showing that reference-copy assumptions are not uniformly valid [50]. Practical follow-up should combine pseudogene-discriminating amplification or long reads, copy-number assessment and family-based phasing where appropriate.

The six-tissue prediction panel is useful for identifying whether model responses are shared or context-dependent, but it should not create physiological roles by inference. The adrenal has direct mechanistic relevance. Normal testis and ovary, adrenal rest tissue, placenta, fetal liver and kidney have different cell populations and hormonal environments. In CAH, gonadal dysfunction may arise from systemic steroid imbalance or ectopic adrenal-like tissue. A similar predicted RNA response across tissues is therefore insufficient to infer a shared local enzyme function. Requirement of POR for CYP21A2 may further complicate simple interpretation, as variations in POR when combined in cytochrome P450 proteins may have differential effects on enzyme activities [51].

The distal 3′ untranslated-region RNA signal is a testable hypothesis emerging from the saturation map. It could motivate reporter assays, endogenous editing and measurements of transcript processing, but the present analysis does not specify which molecular mechanism is responsible. Tissue-appropriate baseline expression should be demonstrated before assigning importance to a predicted fold change in a low-expression context. Where baseline transcript is nearly absent, a regulatory prediction may have little physiological consequence despite a numerical score.

The principal limitations are source coverage, genomic context and calibration. Although the database queries and saturation denominator are auditable, the study is not a manually adjudicated catalogue of every variant in every human genome project or publication. The PubMed search includes a broad inventory and selected primary evidence extraction, rather than a completed systematic review with independent full-text screening. Proprietary mutation databases, unpublished laboratory findings, all population projects and all alternative haplotypes were not exhaustively accessed. ClinVar classifications can change and may include correlated submissions. Population frequencies at this duplicated locus require attention to filters, read mapping and assay design.

The model analysis also uses two contexts: precomputed Atlas scores and direct 16 kb inference. The four control alleles permit inspection of this difference but do not establish general equivalence. The shorter windows exclude distal regulatory context and should not be used for unqualified comparisons with Atlas values. Model gene masks do not exactly equal the MANE transcript used for annotation. Tissue summaries select maxima from unequal track sets, and no patient-specific epigenetic, hormonal or developmental state is represented. Complex structural variants, phase, allele dosage and full conversion haplotypes remain outside the single-allele model.

Priority experiments should be chosen by mechanism and evidence gap. Established splice alleles provide positive controls for endogenous RNA assays; observed intronic variants with substantial usage or junction signals are candidates for splice characterization; coding variants with discordant functional studies merit standardized two-substrate assays with both total-cellular and protein-normalized outputs. Distal untranslated-region candidates require transcript-processing experiments before a causal mechanism is claimed. For all classes, endogenous editing in an appropriate adrenal model would help bridge heterologous assays and native regulation. A successful result should connect a defined allele to an RNA or protein mechanism and then to steroid output, rather than rely on score magnitude alone.

## Conclusions

A reproducible CYP21A2 reference catalogue can combine complete exonic and intronic single-base prediction coverage with observed clinical and population variation. The present resource contains 9,690 Atlas-scored SNVs, 413 additional reported alleles analyzed by direct inference, a reconciled literature inventory and separately preserved functional evidence. Splicing predictions identify mechanisms that total-RNA summaries may miss, while coding pathogenicity often requires biochemical interpretation. Tissue relevance, allele structure and assay normalization remain essential. The catalogue is intended to support variant prioritization, mechanistic experiments and future evidence updates, with explicit boundaries between observation, prediction and clinical inference [52].

## Data availability and reproducibility

The accompanying data are organized into four resources that are available for download at https://doi.org/10.7910/DVN/CZDL3K [53]. Supplementary Data 1 (CYP21A2_variant_catalogue.xlsx) provides the 19-sheet analysis workbook. Supplementary Data 2 (CYP21A2_supplementary_tables.zip) provides the full TSV tables and supporting JSON and RIS files. Supplementary Data 3 (CYP21A2_raw_prediction_scores.zip) provides six compressed score files retaining track-level and junction-level outputs. Supplementary Data 4 comprises CYP21A2_source_data_part1.zip and CYP21A2_source_data_part2.zip, which together preserve the retrieved source records, reference sequences, model metadata, cached predictions and analysis scripts. The two source-data parts retain complementary files within a common folder structure.

The Supplementary data file descriptions identify every workbook sheet and each tabular, score and source-data component. The workbook contains selected columns for convenient inspection; the TSVs and compressed matrices retain fuller annotations and outputs. Original source assertions, missing values and unresolved mappings are preserved. Atlas scores and direct 16,384-bp predictions are labeled separately and should not be pooled without accounting for their different protocols.

The “References” sheet preserves the 37-reference used during analysis bibliography. Its reference_number values are analysis identifiers and do not denote the citation numbers in this manuscript. The manuscript bibliography is more authoritative for its citations. PubMed_inventory.tsv contains the reconciled 1,538-record search inventory, whereas PubMed_article_inventory.ris contains 1,532 journal-article records.

The source-data README documents software requirements, script order limitations and post-processing steps. API credentials are excluded. Scores reflect the API responses obtained at the analysis cutoff of 1 October 2026; future reruns may differ if service models or source databases change.

## Supplementary data file descriptions

**Supplementary Data 1.**
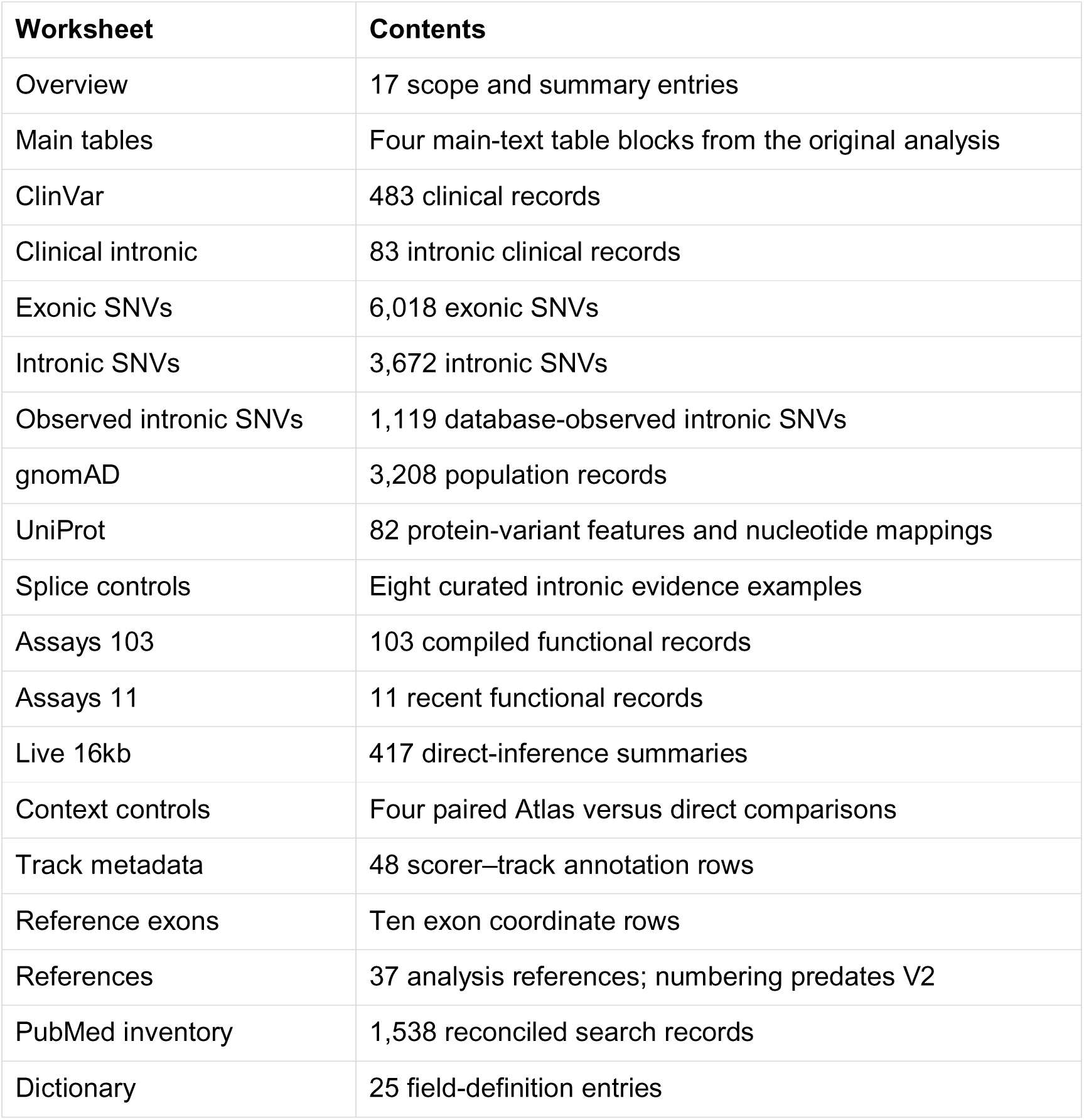
Variant catalogue workbook. CYP21A2_variant_catalogue.xlsx contains the following 19 worksheets. The “Main tables” and “References” sheets preserve the original analysis tables and bibliography; their numeric reference IDs are not the revised manuscript citation numbers.

| Worksheet | Contents |
| --- | --- |
| Overview | 17 scope and summary entries |
| Main tables | Four main-text table blocks from the original analysis |
| ClinVar | 483 clinical records |
| Clinical intronic | 83 intronic clinical records |
| Exonic SNVs | 6,018 exonic SNVs |
| Intronic SNVs | 3,672 intronic SNVs |
| Observed intronic SNVs | 1,119 database-observed intronic SNVs |
| gnomAD | 3,208 population records |
| UniProt | 82 protein-variant features and nucleotide mappings |
| Splice controls | Eight curated intronic evidence examples |
| Assays 103 | 103 compiled functional records |
| Assays 11 | 11 recent functional records |
| Live 16kb | 417 direct-inference summaries |
| Context controls | Four paired Atlas versus direct comparisons |
| Track metadata | 48 scorer–track annotation rows |
| Reference exons | Ten exon coordinate rows |
| References | 37 analysis references; numbering predates V2 |
| PubMed inventory | 1,538 reconciled search records |
| Dictionary | 25 field-definition entries |

**Supplementary Data 2.** Full tables and supporting records. CYP21A2_supplementary_tables.zip contains the files below under output/, together with README.md at the archive root. The full tables preserve additional annotation and source fields beyond the selected workbook columns. main_tables.json and figure_legends.json retain the original analysis presentation records.

| File in output/ | Contents |
| --- | --- |
| exon_reference.tsv | Ten exon coordinates and transcript offsets |
| all_locus_SNV_predictions.tsv | All 9,690 SNVs; full annotations and summaries |
| all_exonic_SNV_predictions.tsv | 6,018 exonic SNVs; includes codon annotation |
| clinvar_catalogue.tsv | 483 clinical records and joined predictions |
| clinvar_intronic.tsv | 83 clinical intronic records |
| gnomad_variants.tsv | 3,208 population records with separate exome/genome data |
| gnomad_intronic.tsv | Intronic population-record subset |
| uniprot_variants.tsv | 82 source protein-variant features |
| uniprot_variants_mapped.tsv | Protein-feature mapping and mapping status |
| PubMed_inventory.tsv | Reconciled 1,538-record inventory |
| PubMed_article_inventory.ris | 1,532 journal articles; six book records excluded |
| published_103_assay_records.tsv | 103 published assay rows with source and quality flags |
| experimental_function_benchmark.tsv | 11-variant assay data and nucleotide mapping |
| track_metadata.tsv | 48 scorer-track annotation rows |
| AVI_SCORE_MODEL_FEATURES.tsv | All 9,690 SNVs and model feature values |
| AVI_SCORE_FEATURE_IMPORTANCE.tsv | All 9,690 SNVs and feature attributions |
| live_16kb_predictions.tsv | 417 direct-inference summaries |
| context_sensitivity_controls.tsv | Four paired context comparisons |
| live_scoring_status.tsv | Completion status for all 417 direct requests |
| known_intronic_SNV_predictions.tsv | 1,119 observed intronic SNVs and source flags |
| intronic_benchmarks.tsv | Eight curated splice-evidence examples |
| statistics.json | Analysis counts and descriptive statistics |
| main_tables.json | Machine-readable original table content |
| figure_legends.json | Original analysis figure legends |

**Supplementary Data 3.**
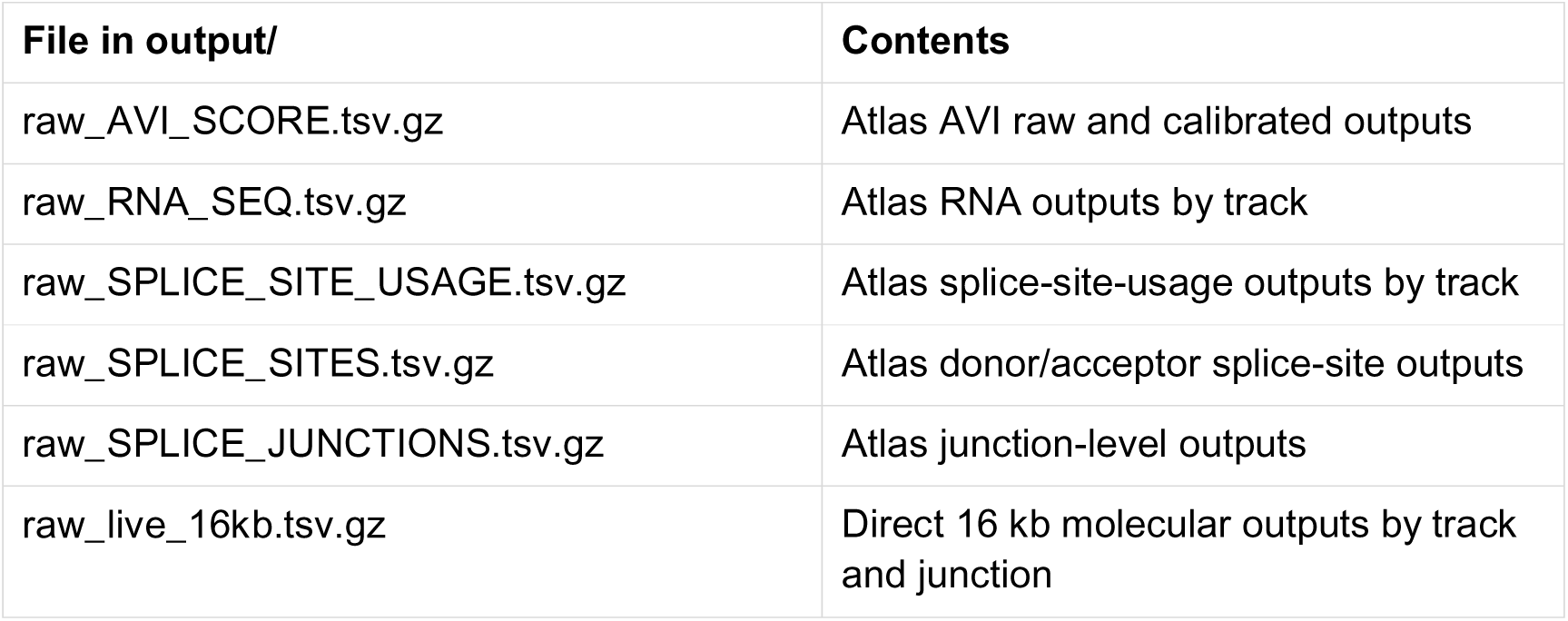
Unaggregated prediction outputs. CYP21A2_raw_prediction_scores.zip contains the following six gzip-compressed TSVs under output/, together with README.md. “Raw” denotes the unaggregated model-score outputs, not experimental sequencing reads. Atlas and direct-inference outputs remain separate.

| File in output/ | Contents |
| --- | --- |
| raw_AVI_SCORE.tsv.gz | Atlas AVI raw and calibrated outputs |
| raw_RNA_SEQ.tsv.gz | Atlas RNA outputs by track |
| raw_SPLICE_SITE_USAGE.tsv.gz | Atlas splice-site-usage outputs by track |
| raw_SPLICE_SITES.tsv.gz | Atlas donor/acceptor splice-site outputs |
| raw_SPLICE_JUNCTIONS.tsv.gz | Atlas junction-level outputs |
| raw_live_16kb.tsv.gz | Direct 16 kb molecular outputs by track and junction |

### Supplementary Data 4 Source data and analysis code

CYP21A2_source_data_part1.zip and CYP21A2_source_data_part2.zip jointly contain the following records. Extract both to a common parent directory while retaining their internal paths; each ZIP can be opened independently. Checkpoint batches are distributed across the two parts and do not constitute different analysis versions.

### Reference and source protein

raw/CYP21A2_region.gb; raw/NM_000500.9.gb; raw/ncbi_gene.json; raw/uniprot.json.

Retrieved genomic and transcript sequences, gene metadata and UniProt source record.

### Clinical retrieval

raw/clinical_search.json; raw/clinical_retrieval_manifest.json; raw/clinical_summary_0.json through raw/clinical_summary_4.json; raw/clinical_vcv_0.xml through raw/clinical_vcv_9.xml; raw/clinical_variants.json; raw/clinical_variants.tsv.

Entrez identifiers, retrieval provenance, five summary batches, ten VCV XML batches and parsed clinical records.

### Population retrieval

raw/gnomad.json; raw/gnomad_filters.json.

Original population payload and filter metadata.

### Literature search and verification

raw/pubmed_full_search.json; raw/pubmed_recovery_audit.json; cyp21a2_pubmed_broad.json; cyp21a2_literature_verified.json; cyp21a2_literature_additional.json; cyp21a2_intronic_additional_verified.json; alphagenome_method_refs.json.

Search and reconciliation records, selected verified bibliography and model-method references; the reconciled inventory is the authoritative final search list.

### Functional extraction

prado2022_103_assays_extracted.json; prado2022_benchmark_tables_verified.json; xu2026_assay_table1_verified.json.

Source table extraction and verification records for the two assay datasets.

### Model metadata

raw/alphagenome_metadata.pkl; raw/atlas_metadata.pkl.

Service metadata retained as cached Python objects.

### Atlas and feature checkpoints

raw/atlas_chunks/*.pkl (26 files); raw/features/*.pkl (13 files).

All locus chunks and feature-query caches; distributed across both archive parts.

### Direct-inference inputs and outputs

raw/known_extra_variants.json; raw/live_16kb/*.pkl (417 files).

Reported non-SNV input representations and individual cached results, including four SNV context controls.

### Analysis code and documentation

scripts/run_atlas.py; scripts/run_features.py; scripts/run_known.py; scripts/run_live.py; scripts/analyze.py; scripts/integrate.py; scripts/aggregate_live.py; scripts/figures.py; scripts/manuscript.py; scripts/prepare_workbook.py; scripts/workbook.mjs; requirements.txt; README.md; SHA256SUMS.txt.

Request, annotation, integration and presentation scripts, environment requirements, workflow caveats and original content checksums. run_known.py records the initial unsupported Atlas non-SNV route; run_live.py supplies the successful direct protocol.

The raw/atlas_chunks/ and raw/features/ folders contain 26 and 13 cached files, respectively; raw/live_16kb/ contains 417 cached results. The part-wise counts are 12/14 Atlas chunks, 5/8 feature files and 209/208 direct-result files. The metadata files comprise one cached object in each part. README.md is repeated for convenience. SHA256SUMS.txt describes the original combined archive contents and is intended for verification after the components have been reunited.

## Declarations

This analysis used publicly available aggregate genetic, bibliographic and functional data. No new patient samples, identifiable individual data or animal experiments were analyzed. Clinical classifications were reproduced as database assertions and were not independently reassigned.

